# Shifting human-rodent interfaces under climate change: modeling the distribution of the reservoir for Junin virus and associated drivers

**DOI:** 10.1101/2024.06.24.600371

**Authors:** Nuri Flores-Pérez, Pranav Kulkarni, Marcela Uhart, Pranav Pandit

## Abstract

**Background:** The drylands vesper mouse (*Calomys musculinus)* is the principal host for *Junin mammarenavirus* (JUNV), which causes Argentine Hemorrhagic Fever (AHF) in humans. In our study, we aimed to assess the probable range of *C. musculinus* and to identify hotspots for potential disease transmission to humans under current and future climate change scenarios.

**Methodology/Principal Findings:** We used tree-based machine learning (ML) classification algorithms to generate and project *C. musculinus* habitat suitability under two climate change scenarios for the years 2050 and 2070 using bioclimatic and landscape related predictors. Evaluation of the models showed high accuracy, with AUC_ROC_ ranging from 86.81-89.84%. The analysis of the importance and influence of different variables indicated that the rodent prefers warm temperatures, moderate annual precipitation, low precipitation variability, and low pasture coverage. While a severe climate change scenario (Representative Concentration Pathway 8.5) suggests a reduction in suitable areas for JUNV reservoir and a decrease in hotspots for potential disease transmission, an intermediate scenario (Representative Concentration Pathway 4.5) displays expansion in areas for *C. musculinus* distribution alongside increased potential hotspot zones.

**Conclusions/Significance:** While acknowledging the complexity of ecological systems and the limitations of the species distribution models, our findings offer a framework for developing preventive measures and conducting ecological studies in regions prone to *C. musculinus* expansion and hotspots for potential disease transmission driven by climate change. Preventive interventions will need to be adapted to target *C. musculinus* changing spatial dynamics.

**Author Summary:** Climate change might modify where animals live, including those that carry diseases than can spread to humans. This study focused on how climate change might affect the drylands vesper mouse (*Calomys musculinus*). This rodent carries the Junin virus, which causes Argentine Hemorrhagic Fever, a dangerous disease if left untreated. We used species distribution models to predict how this rodent’s range might change by 2050 and 2070 under an intermediate and severe climate change scenario.

We found that a temperature rise of 1.1°C to 2.6°C (intermediate change scenario) could allow the rodent to move into new areas, potentially increasing disease transmission to humans. These new areas might have warm temperatures, moderate rainfall, and low grass cover, as we found that *C. musculinus* prefers those climate and landscape conditions.

Knowing the rodent’s potential distribution and preferred habitat enables preparedness and monitoring for early warning systems. By establishing monitor systems and online platforms featuring species distribution models and predictions, we can facilitate communication and resource sharing among researchers, policymakers, healthcare professionals and the public. These tools will help protect public health as climate change continues to bring new challenges.

## Introduction

Anthropogenic climate change has caused an increase in global temperatures. By 2022, the average global temperature was about 1.15°C above the 1850-1900 average [1]. It is predicted that in the next two decades, global warming will continue to increase, potentially causing more frequent heatwaves and droughts, and more intense and frequent heavy precipitation [2]. Furthermore, there will be long periods of drought in sub- tropical and mid-latitudes, interspersed with intense rainfall events [2].

The effect of these global changes needs to be considered in the study of zoonotic infectious diseases, as hosts and vector species will likely experience shifts in their assemblages, migrate to new areas, or modify their abundance. This may increase overlap with humans, facilitating the transmission of diseases. For example, in South American countries and the United States, an increase in food availability for peri- domestic rodents after intense rainfall and flooding caused outbreaks of Hantavirus Pulmonary Syndrome [4,5]. Viral spillover events into human populations have been associated with wildlife movements following droughts or wildfires [6]. Importantly, human migration and urbanization in response to climate change may also influence disease risk as areas with high human population density are predictive of infectious disease emergence [7,8].

Species distribution models (SDM) have been used to model the potential distribution of animal or plant species based on observed occurrences and a set of measured or estimated environmental variables [9]. As the predicted range shifts vary according to taxonomic groups, detailed information about species’ physiological, ecological, and environmental data is needed to make specific predictions for individual species [10]. The inclusion of climate change scenarios to evaluate shifts in the distribution of species is relevant and has been frequently used to predict species habitat suitability [11,12]. For example, an environmental-mechanistic model developed by Redding et al. [13] revealed that Lassa fever virus (LAS) is likely to spread to more areas in West Africa, where hotter temperatures and wetter conditions could provide a more suitable habitat for the LAS virus reservoir, the natal multimammate mouse (*Mastomys natalensis*).

In South America, previous studies have used the maximum likelihood method [14] and cellular automaton models [15] to predict the potential distribution of *Calomys musculinus* (*Cricetidae*: *Sigmodontinae*) using bioclimatic and landscape data. This rodent is found in the countries of Argentina, Bolivia, and Paraguay. It occupies a wide variety of habitats, like natural grasslands, shrub steppes, crop field borders, and human-disturbed peri- domestic environments such as wastelands, railroads, and urban garbage dumps [16,17]. The abundance of *C. musculinus* changes seasonally, driven by natural climatic variations and land-use practices that affect resource availability [18]. Moreover, corn crops and land-surface temperature are positive predictors of the rodents’ abundance. Other predictors such as crop field borders offer habitat, food, and shelter year-round [19].

*C. musculinus* is the main reservoir host of the *Junin mammarenavirus* (JUNV) [20,21], the etiological agent of Argentine Hemorrhagic Fever (AHF). This disease is endemic to the humid pampas in Argentina, a densely populated area at the core of the largest agro- industrial complex in the country [22,23]. In severe cases of AHF, patients develop hemorrhagic and neurological complications, with a fatality rate of 20% [24]. AHF was first reported in 1955 among agricultural workers [20]. It is believed that the outbreak was due to land use changes and occupational exposure via agricultural activities [25]. Since its emergence, a constant and progressive extension of the AHF endemic area has been observed [26]. In the last 10 years, the disease has expanded to northeastern Argentina, with reemergence in areas that had not had cases for 15-20 years [27].

In 1992, the use of Candid#1 vaccine was implemented, causing a decrease in AHF incidence [28, 22]. However, in the following years, changes in the risk patterns were observed, with increased cases among women, children under 15 years old, and people who work in non-rural jobs and/ or reside in urban zones [22].

Understanding how climate change may influence the distribution of *C. musculinus* will help assess the spatial risk factors for AHF transmission. This information will be crucial for public health surveillance and for guiding preventive interventions. In this study, we developed a SDM based on bioclimatic and landscape variables for *C. musculinus*. We incorporated machine learning (ML) classifiers and considered future climate change scenarios to anticipate potential shifts in the rodent spatial distribution to address three objectives: (i) to evaluate the importance and impact of different bioclimatic and landscape variables in the prediction of *C. musculinus* range shifts, (ii) to assess these geographic changes under current and future climate change scenarios, and (iii) to identify hotspots for potential disease transmission.

## Methods

### Study area

The study was focused on Argentina (WGS 84 bounds: -73.6°, -53.7°, -55.1°, -21.8°) to assess the current and future habitat suitability for *C. musculinus* across diverse geographical and ecological regions within the country. Argentina, covering a land area of 2,780,400 km^2^ and 3,694 km north-south span, has fertile lands, mainly in the central and the eastern pampas plains and irrigated valleys at the base of the Andes Mountains [29]. Agriculture is one of its principal economic activities, consisting of monocultures of corn, soybean, sunflower, and wheat [18]. The human population is predicted to grow from the current 46 million to 73.4 million by 2100 [30,31].

Argentina can be divided into 18 ecoregions, which are territories with relatively uniform or recurrent environmental conditions [32] (S1 Fig). Notably, there has been a progressive extension of the AHF endemic region into the north-central area of the Pampas ecoregion. This expansion poses a potential risk to approximately 5 million people [21].

### Species occurrence data

Presence data for *C. musculinus* was retrieved from the Global Biodiversity Information Facility (GBIF; [33]), limiting the occurrences to the bounds of Argentina. To align the presence data period closely with the bioclimatic and landscape variables period, we constrained the dataset to records spanning from 1990 to the last available year of *C. musculinus* records. After deleting duplicates and missing values, the refined dataset consisted of 70 observations from 1990 to 2021 (Fig 1).

**Fig 1.**
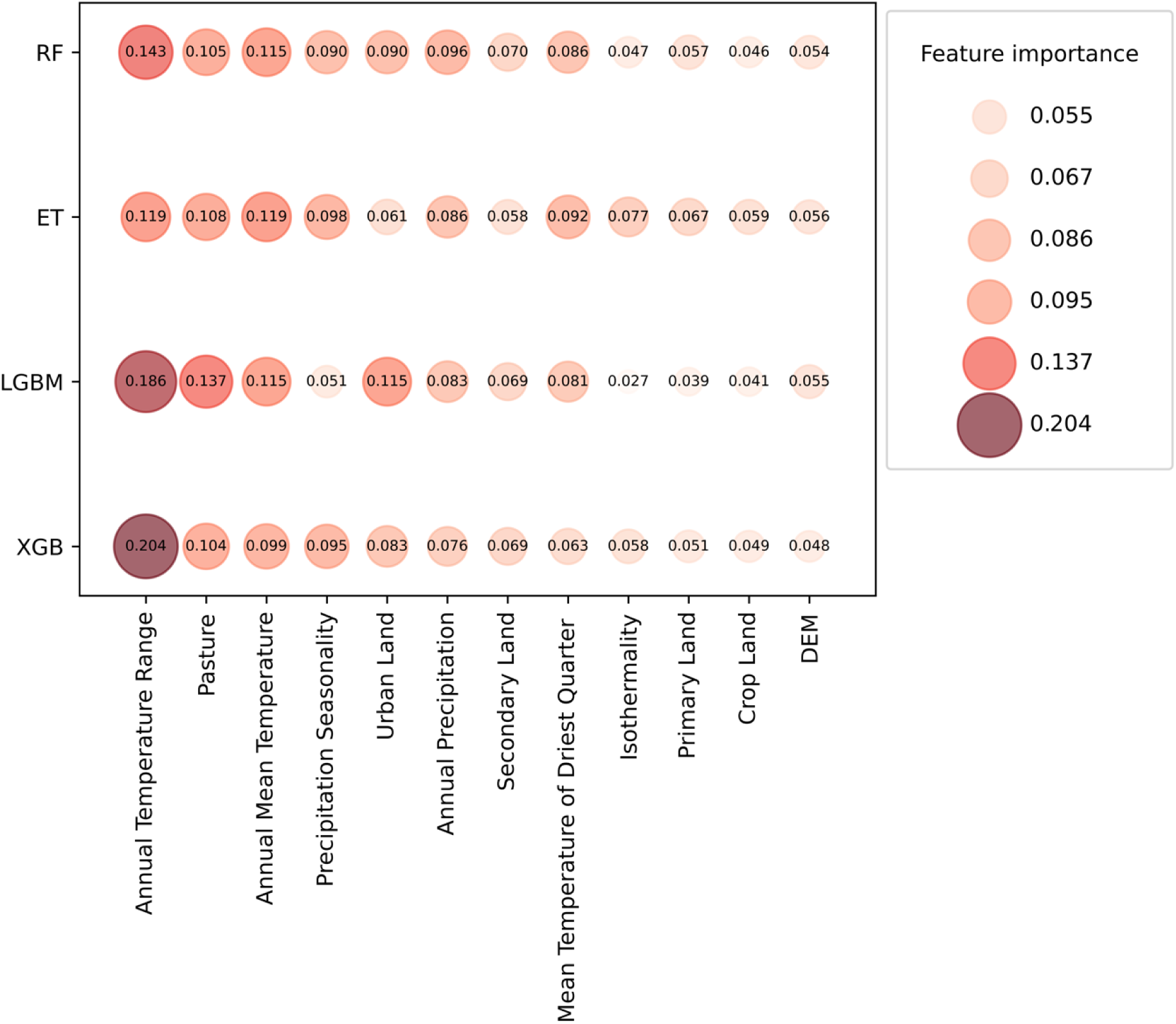
Importance of bioclimatic and landscape variables used for predicting *C. musculinus* distribution. The influence of bioclimatic and landscape variables is shown across the four classifiers used for predicting AHF reservoir distribution: Random Forest (RF), Extra Trees (ET), Extreme Gradient Boosting (XGB), and Light Gradient-Boosting (LGBM). Note: Bigger circles and higher values indicate a greater importance.

### Predictor bioclimatic and landscape variables

We selected 26 bioclimatic and landscape variables (S1 Table) as predictors for our species distribution model, based on relevance to *C. musculinu*s ecology and data availability. Of these 26 variables, 19 were bioclimatic variables obtained from WorldClim [34], five were land use variables (percentage of cropland, primary and secondary land, pasture, and urban land) retrieved from Harmonized Global Land Use [35], and one was an elevation variable from the Consortium for Spatial Information [36]. Because the feature importance assessment (explained below) can neglect the importance of correlated features [37], we utilized the Pearson correlation coefficient to exclude highly correlated variables (r ≥ 0.7) and built our model with only non-correlated features (Table 1).

**Table 1.**
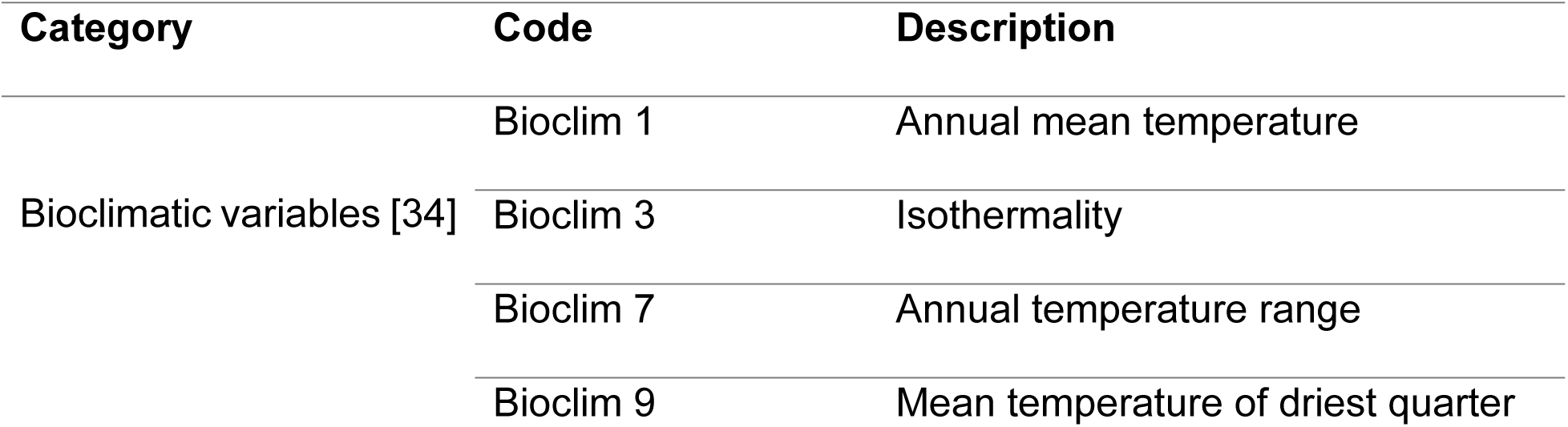
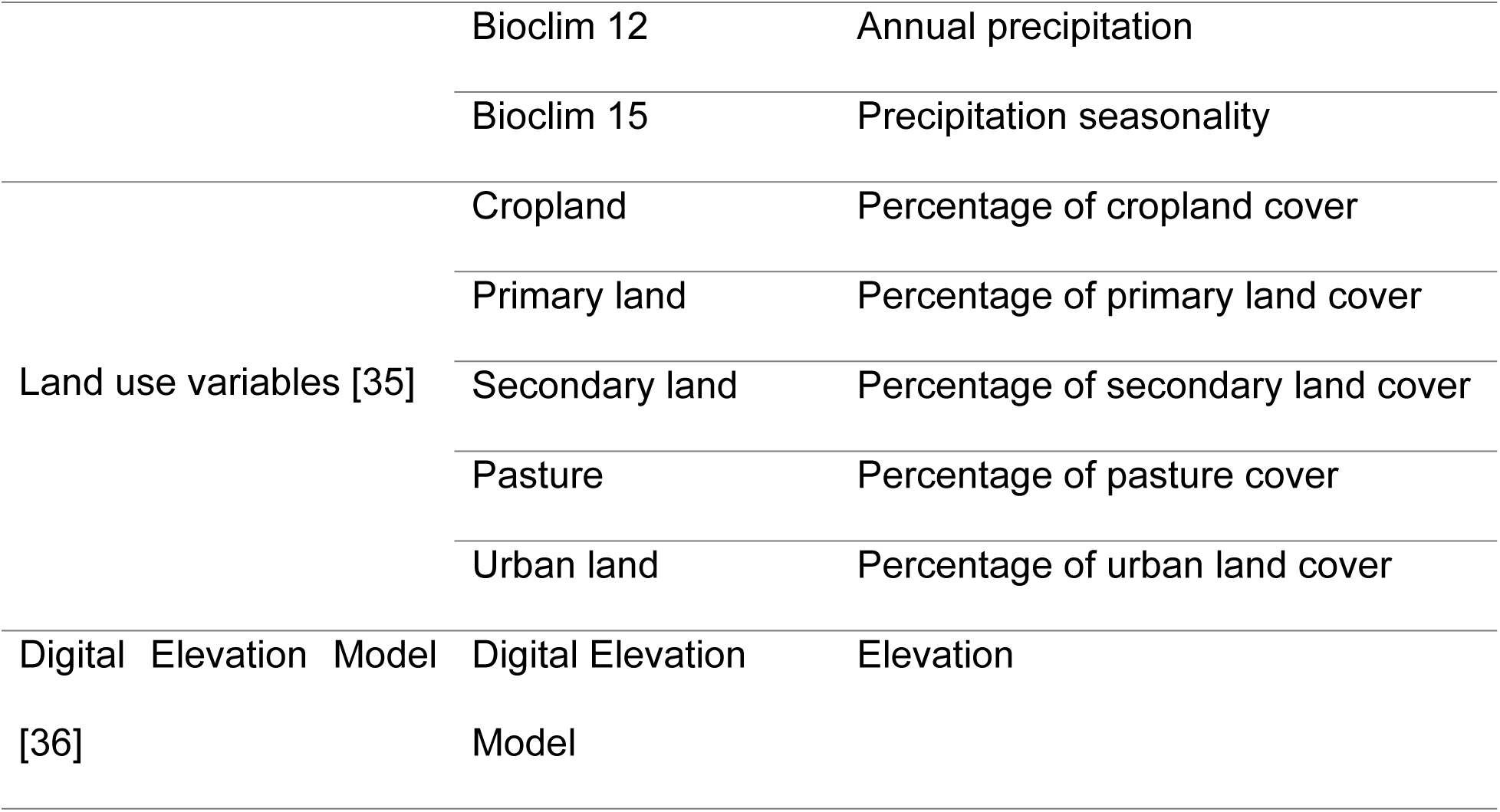
Bioclimatic and landscape variables used to build *C. musculinus* species distribution models.

To calculate the predicted habitat suitability in future climate change scenarios, we used two Representative Concentration Pathways (RCPs) from the Coupled Model Intercomparison Project Phase 5 (CMIP5; [38]): RCP4.5 (intermediate greenhouse gas emissions) and RCP8.5 (high greenhouse gas emission) for years 2050 and 2070. RCPs represent a range of possible future radiative forcing levels based on emissions of greenhouse gases and aerosols [39].

All raster variables were sourced to GeoTiff files and resampled to align with the dimensions (597 x 999), resolution (0.042 degrees), and projection (WGS84) of the bioclimatic variables. This ensured consistent spatial representation across all raster files, facilitating accurate integration for the species distribution model analysis.

### Species distribution models

To ensure robustness and reliability in the modeling process and results, we generated 1000 training datasets, each with 70 presence points paired with 70 randomly generated pseudo-absences points. We created pseudo-absences with the *randomPoints()* function in R software version 4.2.2; using a 1:1 ratio between presence and pseudo-absences yielded the highest accuracy in our predictive models, as described by Barbet-Massin et al. [40].

We employed an ensemble approach to build the potential *C. musculinus* distribution, utilizing four Machine Learning (ML) algorithms: Random Forest (RF), Extra Trees (ET), Extreme Gradient Boosting (XGB), and Light Gradient-Boosting (LGBM). ML models were trained in Python version 3.10.9 using the scikit-learn library [41] for the RF and ET algorithms. For the XGB algorithm, we utilized the XGBoost library [42], and for the LGBM algorithm, the LightGBM library [43]. For all ML algorithms, the default hyper-parameters were used during the training of classifier models. Model predictions for current and projected scenarios were generated using the *impute* function from the PyImpute library in Python 3.8 [44]. Each ML classifier model was trained 1000 times with randomly generated pseudo-absences and predictions were averaged from each classifier to generate habitat suitability maps for *C. musculinus*. For each iteration of the model, we evaluated the algorithm’s performance through 5-fold cross-validation using three measures of predictive performance: (i) area under the ROC curve (AUC_ROC_), (ii) recall and (iii) precision.

Similarly, for each iteration, the importance of each variable was evaluated using the Mean Decreased Impurity (MDI) for the RF and ET classifiers [45]. For XGB and LGBM classifiers the Gain Feature Importance was calculated [46]. For each ML classifier, we averaged the 1000 simulation results and generated boxplots to visually summarize the findings. We generated Partial Dependence Plots (PDPs) to understand how changes in the values of the input variables influence the target response (presence of the species) [47]. To generate the PDPs, we also averaged each classifier’s 1000 simulation results. Finally, to determine the predicted change in *C. musculinus* distribution — whether it would expand, contract, or remain the same — we used the following formula for each scenario:

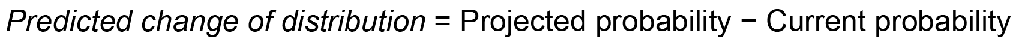

### Hotspots for potential disease transmission

We obtained Argentina’s human population density data from the Socioeconomic Data and Applications Center [48]. This information is based on the Shared Socioeconomic Pathways, consisting of global population data for the base year 2000 and projections at ten-year intervals for 2010 to 2100 [49].

We set a probability threshold greater than 60% (p > 0.6) for the presence of *C. musculinus*, under the assumption that areas surpassing this threshold could exhibit habitat suitability conducive to sustaining a high rodent population density, thus supporting the transmission of JUNV [26]. Regarding the human population density, we established three tiers: (i) human population density greater than 1000, (ii) 10,000 and (iii) 100,000. These thresholds were chosen to capture areas with varying human population densities, ensuring that hotspots are identified in regions with potential human-rodent overlap and subsequent disease transmission. We generated a raster layer to represent areas that satisfied both criteria, indicating regions where the threshold of *C. musculinus* presence probability overlapped with the human population density thresholds. These areas were classified as hotspots, delineating zones of potential disease transmission.

## Results

### Model comparison and evaluation

After omitting 13 highly correlated bioclim variables (r ≥ 0.7) (S3 Fig) our model included the following variables: bioclim 1 (Annual Mean Temperature), bioclim 3 (Isothermality), bioclim 7 (Annual Temperature Range), bioclim 9 (Mean Temperature of Driest Quarter), bioclim 12 (Annual Precipitation) and bioclim 15 (Precipitation Seasonality), all land use variables, and elevation (DEM) (Table 1).

Table 2 shows the simulation’s average results of AUC_ROC_, precision, and recall scores for four tree-based ML algorithms used to predict *C. musculinus* presence. The algorithm with the highest AUC_ROC_, precision, and recall was Extra Trees (ET) followed closely by Random Forest (RF). On the other hand, Light Gradient-Boosting (LGBM) had the lowest scores across all three metrics.

**Table 2.** Performance results of the machine learning classifiers used for predicting ***C. musculinus* distribution.**

| Machine learning algorithm | $AUC_{ROC}$ | Precision | Recall |
| --- | --- | --- | --- |
| Random Forest | 89.57 (+/- 10.35) | 80.64 (+/- 18.55) | 84.54 (+/- 18.40) |
| Extra Trees | 89.84 (+/- 9.74) | 80.99 (+/- 18.52) | 85.30 (+/- 17.71) |
| Extreme Gradient Boosting | 87.44 (+/- 11.51) | 78.75 (+/- 19.14) | 82.41 (+/- 19.16) |
| Light gradient-boosting | 86.81 (+/- 11.77) | 78.19 (+/- 19.18) | 82.36 (+/- 19.40) |

### Importance of bioclimatic and landscape variables for *C. musculinus* distribution

The influence of each variable across the four algorithms is presented in Fig 1, while a summary of the 1000 simulation results of feature importance can be found in the supplementary material (S2 Fig). For RF, the most important feature was the annual temperature range, followed by the annual mean temperature and the pasture feature. Similarly, the annual mean temperature ranked higher for ET, followed by the annual temperature range and the pasture feature. For both XGB and LGBM, the most important feature was the annual temperature range, followed by the pasture feature. The XGB classifier had the annual mean temperature in the third place, while the LGBM had urban land.

Fig 2 shows single-variable partial dependence plots for key bioclimatic and landscape variables across classifiers. The species suitability increased when the annual temperature range increased, but it decreased when the temperature reached 30°C, according to the ET classifier. In the RF, XGB and LGBM, the response started weak, increasing after 20°C and decreasing around 30°C. For the annual mean temperature, in all the classifiers, after 5°C there was increasing suitability when the temperature increased, after that, there was a decrease around 17°C. As for the annual precipitation, there was a weak response until 700 mm of total annual precipitation, where the species’ partial dependence started to decrease as the precipitation increased. On the other hand, the rodent’s suitability increased as the precipitation seasonality increased, but the response appeared weak at around 80% of precipitation variability, except in the ET classifier, which showed a decrease in suitability as precipitation seasonality increased. For the RF, XGB, and LGBM classifiers, the partial dependence of *C. musculinus* presence increased with increasing pasture coverage until 10% coverage. The partial dependence of the rodent decreased by around 20% for all ML models, becoming weak after 50%. In particular, the ET classifier showed a constant decrease in partial dependence after 20% of pasture cover. For the urban land feature, the rodent’s presence increased with increasing urban land coverage but became weak over 0.5% of urban land coverage.

**Fig 2.**
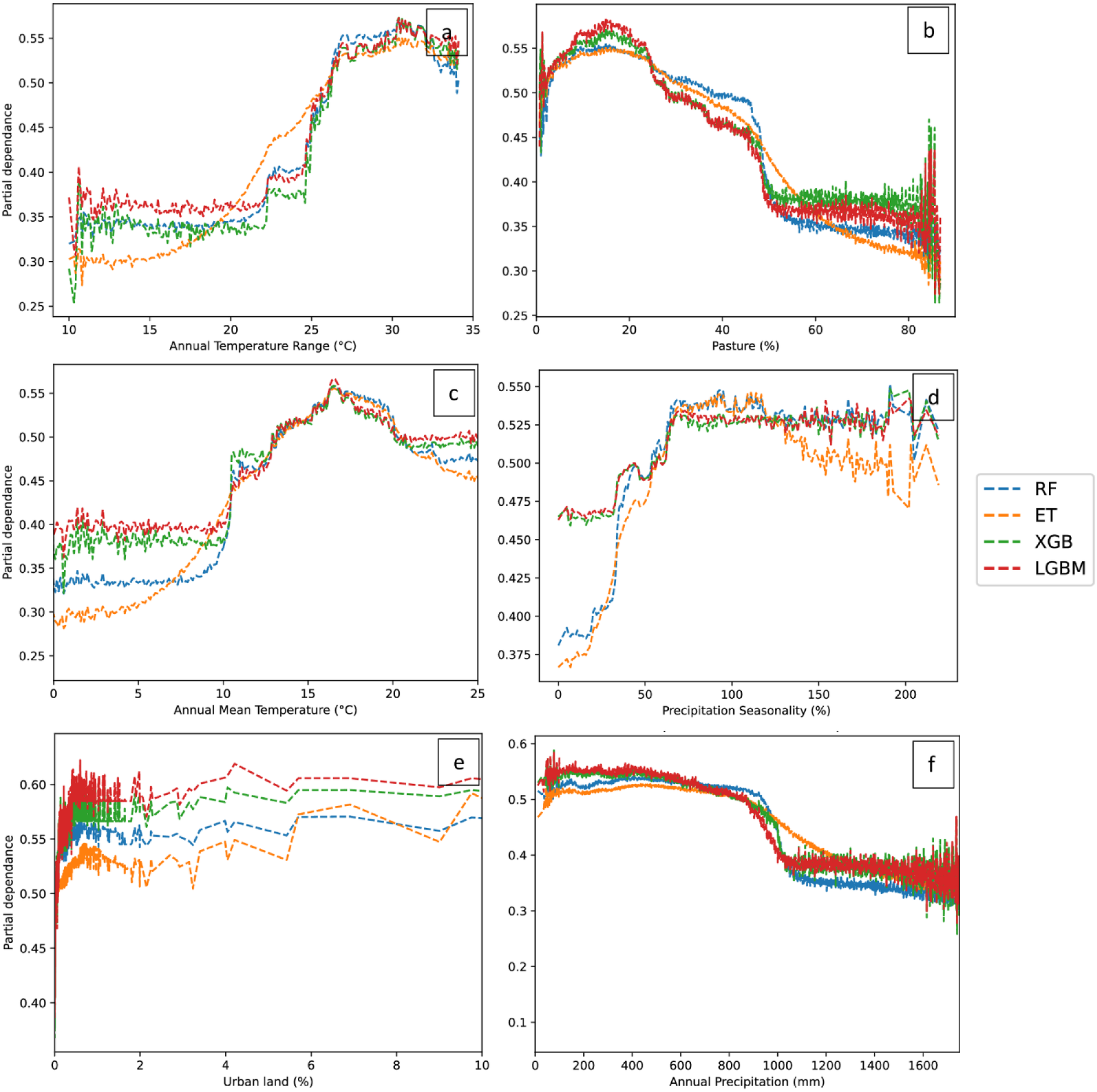
Partial dependence of *C. musculinus* distribution on key bioclimatic and landscape variables. a) Annual temperature range (°C), b) Pasture (percentages of pasture coverage), c) Annual mean temperature (°C), d) Precipitation seasonality (percentage of precipitation variability), e) Urban land (percentages of urban land coverage), f) Annual precipitation (millimeters of rainfall). Note: The color of the lines indicates the machine learning classifier used: Random Forest (RF) in blue, Extra Trees (ET) in orange, Extreme Gradient Boosting (XGB) in green and Light Gradient-Boosting (LGBM) in red.

### Distribution of *C. musculinus* under current and climate change scenarios

The *C. musculinus* current distribution prediction matched *C. musculinus* occurrence data from GBIF and showed a high probability of occurrence in the Pampas, Espinal, Monte de Llanuras y Mesetas, Chaco seco, Monte de Sierras y Bolsones, and Yungas ecoregions (Fig 3).

**Fig 3.**
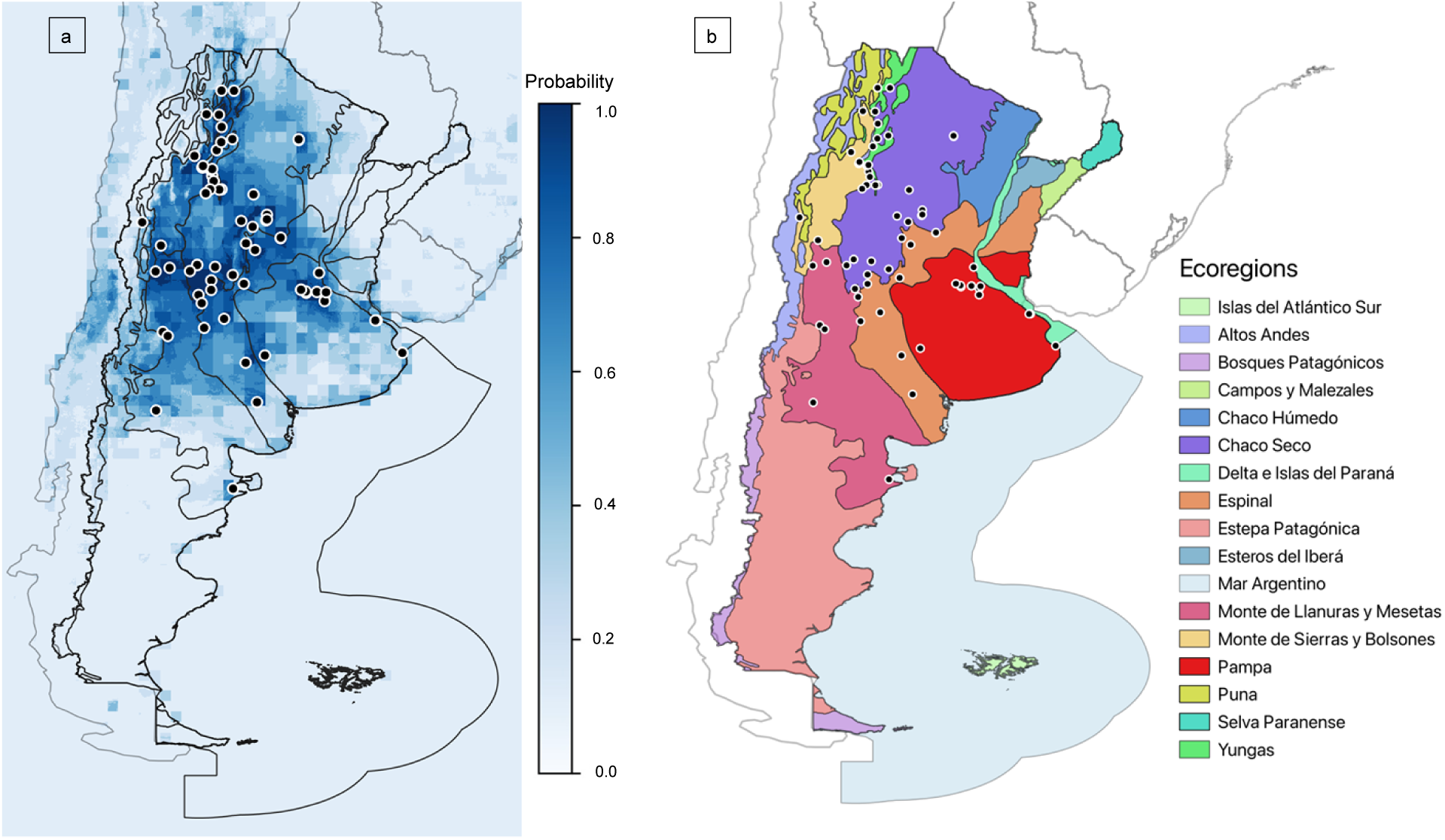
Predicted distribution of *C. musculinus* in Argentina’s ecoregions under current bioclimatic and landscape conditions. a) Current predicted distribution of *C. musculinus* with higher values and darker blue colors indicating a greater likelihood of presence. b) Argentina’s ecoregions as defined in Methods section, overlaid with *C. musculinus* occurrence data from GBIF (black dots).

Fig 4 displays the predicted change in *C. musculinus* distribution — whether it would expand, contract, or remain the same between the current conditions and under the RCP4.5 and RCP8.5 climate change scenarios. *C. musculinus* distribution under the RCP4.5 scenario for 2050 and 2070 mainly showed contraction at the northeast of the Pampas ecoregion, northern Espinal, south of Chaco Seco and Monte de Sierras y Bolsones, the northern area of Monte de Llanuras y Mesetas, and most areas of the Yungas and Selva Paranaense. The rest of the ecoregions showed areas of expansion, and there were a few areas with no change in *C. musculinus* distribution, such as the southern area of Estepa Patagónica ecoregion. The predicted areas of expansion or contraction in both scenarios are similar, but in 2070 they showed a decrease in expansion and an increase in contraction.

**Fig 4.**
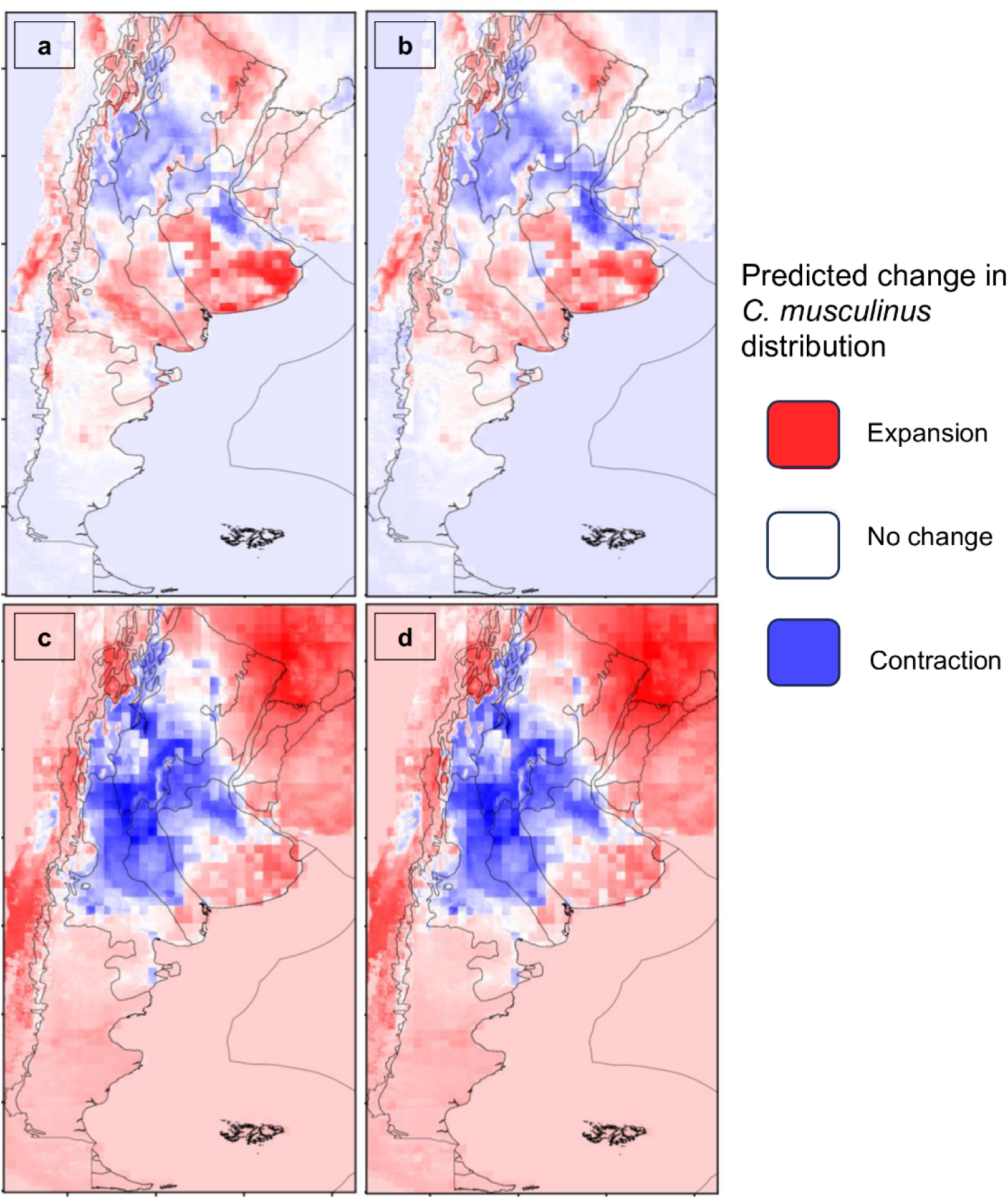
Predicted change in *C. musculinus* distribution between current and future climate change scenarios. Expansion (red), contraction (blue), and areas with no predicted change (white) between current and future climate change scenarios and years: RCP4.5 for year 2050, b) RCP4.5 for year 2070, c) RCP 8.5 for year 2050, and d) RCP 8.5 for year 2070.

In the RCP8.5 scenario for 2050 and 2070, there were areas of expansion in the east Espinal, Esteros del Iberá, Campos y Malezales, Selva Paranaense, Chaco Húmedo, Bosques Patagónicos, Altos Andes, Puna, the south of Estepa Patagónica, the southern Pampas, the north of Chaco seco, south of Espinal, and the north of Yungas. The rest of the ecoregions presented contraction in *C. musculinus* distribution. Between 2050 and 2070, the expansion increased, especially in the north and south. On the other hand, the contraction increased in Monte de Llanuras y Mesetas and Monte de Sierras y Bolsones. Between RCP4.5 and RCP8.5 for both 2050 and 2070, the predicted contraction in the RCP8.5 scenario extended and increased in Monte de Llanuras y Mesetas, Monte de

Sierras y Bolsones, Espinal, the west Pampas and most areas of Chaco seco. Moreover, the predicted expansion increased in the north and south of Argentina, also reaching neighboring Chile, Paraguay, Brazil, and Bolivia.

### Hotspots for potential disease transmission

Hotspots for potential disease transmission can be seen in Fig 5. In the current conditions and the two climate change scenarios, there was a decrease in hotspots when the human population density increased. Under current conditions, there was a concentration of hotspots in the Pampas, Espinal, Chaco Seco, Yungas, Monte de Llanuras y Mesetas, and Monte de Sierras y Bolsones ecoregions. In the RCP4.5 scenario, hotspots were present in the same ecoregions but their size increased. In 2070, the number of hotspots decreased compared to 2050. On the other hand, in the RCP8.5 scenario for the year 2050, there were a couple of hotspot areas in Pampas, Altos Andes, Monte de Sierras y Bolsones, Yungas and Puna. The year 2070 in the RCP8.5 scenario displayed hotspots in Pampas, Altos Andes, Puna, and Chaco Húmedo. For this latter year and scenario, there were no hotspots for the 100,000 human density population threshold.

**Fig 5.**
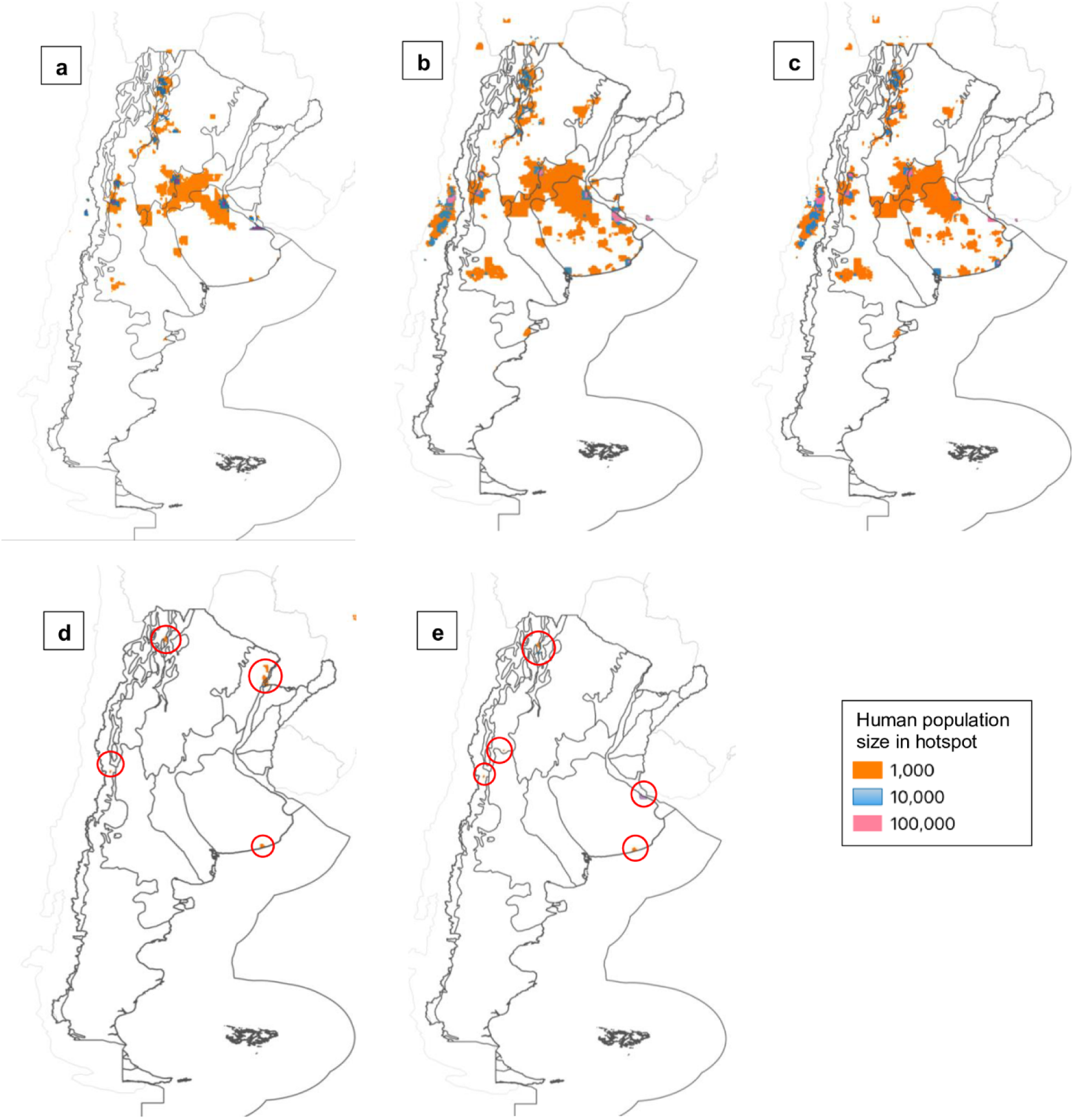
Hotspots for potential AHF disease transmission for current and future climate change scenarios. Hotspots are displayed according to three total human population density thresholds (1,000, 10,000 and 100,000) under: a) current conditions, RCP4.5 scenario for year 2050, c) RCP4.5 scenario for year 2070, d) RCP8.5 scenario for year 2050, and e) RCP8.5 scenario for year 2070. Note: Red circles point out the predicted hotspots in the RCP8.5 scenario.

## Discussion

In this study, we explored the potential distribution of *C. musculinus*, the principal reservoir host for the JUNV, under current and future climate change conditions. We also identified bioclimatic and landscape drivers better describing the changes in the ecological niche of the reservoir rodent. Additionally, we predicted hotspots for potential AHF transmission. Our findings indicated contrasting trends: in the severe climate change scenario (RCP8.5) there were greater contraction areas than in the intermediate scenario (RCP4.5). Also, the latter scenario had more hotspots for potential disease transmission when compared to the RCP8.5 scenario and current climate conditions. Furthermore, our research revealed insights into the habitat preferences of *C. musculinus,* indicating a tendency towards environments with warm temperatures, moderate annual precipitation, and a low precipitation variability, as well as low pasture coverage. Our models complement previous studies [14,15] by providing insights into how climate change might impact the distribution of *C. musculinus* and by highlighting areas of potential disease transmission to humans.

For current climate conditions, our species distribution model aligned with the presence locations of C*. musculinus* from the GBIF database. The predicted distribution also matched the distribution of the endemic area for AHF. Moreover, it aligned with the most recent assessment for *C. musculinus* [50] which reports presence in the following ecoregions: Altos Andes, Puna, Yungas, Chaco Seco, Delta e Islas del Paraná, Espinal, Pampas, Monte de Sierras y Bolsones, Monte de Llanuras y Mesetas, and Estepa Patagónica. The predictions under the two RCP scenarios for the years 2050 and 2070 provided important insights into the influence of climate change on the habitat suitability for the AHF rodent reservoir. The observed areas of expansion in both scenarios signaled a potential shift in the species’ range, yet there was a discernible decline in the predicted expansion in the RCP4.5 scenario for the year 2070 versus the year 2050. This suggests that while certain regions may initially offer habitat suitability for the species, these areas may not be sustainable in the long term. Conversely, in the RCP8.5 scenario the predicted expansion intensified in the northern and southern areas of Argentina when comparing the year 2050 vs 2070, suggesting that for this scenario and in those areas, the bioclimatic and landscape conditions may become increasingly favorable for the species over time. Furthermore, when contrasting both climate change scenarios, we observed a larger area of contraction in the RCP8.5 scenario. This could indicate that under a severe climate change scenario, there will be less favorable bioclimatic and landscape conditions for *C. musculinus* in most of the country’s ecoregions. Nevertheless, in this scenario, the predicted expansion increased in northern and southern Argentina, as well as surrounding countries, such as Chile, Paraguay, Brazil, and Bolivia.

As regions become more suitable for *C. musculinus* in certain areas, there may be increased dispersal and migration, which could facilitate the spread of JUNV to new areas. Nonetheless, other conditions must exist for viral transmission, such as high *C. musculinus* population abundance and communities with low diversity of rodent species [51, 26]. If areas with these conditions exist, there could be an elevated risk of AHF transmission to humans, if human population density is also high. Our results indicated that under a severe climate change scenario (RCP8.5), there would be fewer hotspots for potential disease transmission. Conversely, an intermediate scenario (RCP4.5) suggested an increase in hotspot areas when compared to the current bioclimatic and landscape conditions and the severe climate change scenario. Consequently, public health interventions in Argentina aimed at reducing the risk of JUNV transmission may need to be adapted to these changing spatial dynamics and focus on areas with persistent predicted hotspots in the future.

Our findings, coupled with previous studies, suggest that the expansion of *C. musculinus* may be influenced by (a) warmer temperatures, particularly during fall and winter, (b) precipitation patterns conducive to resource availability, and (c) land use changes, such as reduced pasture coverage, which could favor the abundance of *C. musculinus*. These factors emerged as the critical drivers for the rodent’s potential expansion into regions that will be characterized by these conditions in the future. Moreover, regions projected to experience bioclimatic and landscape changes aligning with these preferences could become hotspots for potential AHF transmission, particularly in areas with high human population density.

The annual temperature range, annual mean temperature, annual precipitation, precipitation seasonality, pasture and urban land coverage were important factors in improving the decision quality of the ML classifiers used in the study. As for the bioclimatic variables, the results suggested that *C. musculinus* prefers annual mean temperatures between 10-17°C, and annual temperature ranges of 22-27°C. This coincides with Simone et al. [19] who documented a positive correlation between rodent abundance and land-surface temperatures during the coldest periods. Furthermore, an annual precipitation greater than 800 mm of rainfall and precipitation seasonality greater than 70% may decrease the habitat suitability for the rodent. These results align with previous studies which have demonstrated that the role of precipitation in rodent abundance varies according to the season, where high precipitation in spring and summer favors population growth by increasing plant resources but causes higher mortality in winter [52, 53].

The analysis of the land cover features revealed that the habitat suitability for *C. musculinus* decreased when pasture coverage reached 20%. A possible theory for this might be that higher pasture coverage areas are dominated by specialist rodent species, such as Azara’s grass mouse (*Akodon azarae)*, which can displace *C. musculinus* to areas with less vegetation cover [54]. Furthermore, the urban land feature appeared to improve the decision quality of the LGBM and XGB classifiers. The corresponding partial dependence plot showed that the presence of the species increased as urban land coverage increased, but after 0.5% coverage the response flattened (asymptotic). The predictive power of our models depends on the validity of extrapolation, which emphasizes the importance of a cautious approach to interpretation. However, the initial increase could indicate that peri-urban zones might indeed be suitable for the species, aligning with Castillo et al. [55], who observed abundance in unoccupied areas, garbage dumps, and railway and stream edges within the city of Río Cuarto, Córdoba, Argentina. Nevertheless, Chiappero et al. [56] suggested that these urban rodent populations might have originally been native to the area, but subsequently encircled by the city’s expansion. If the encircling of rodent habitats by urbanization continues, it becomes necessary to examine the potential implications for public health. For example, it has been observed that the risk of Hantavirus Pulmonary Syndrome (HCPS) increases in areas where human activities, such as urbanization and habitat fragmentation, encroach on rodent habitats, as seen with infected rodents in Uruguay. Similar patterns are observed with other zoonotic diseases like Nipah and Ebola viruses [57]. Understanding these dynamics will be crucial for developing effective public health strategies and mitigating risks associated with rodent-borne disease.

Although previous studies have found a correlation between crops/ crop borders and the presence of *C. musculinus* [19, 58], this variable was not among the most important features in our model. One possible explanation is that the rodent occurrence data was recorded more frequently in pasture lands rather than croplands, thereby adding sampling bias to our models. Despite this result, it is recognized that climate change can lead to modifications in production and agricultural practices [59]. In Argentina, warmer and wetter conditions could cause an extension of croplands towards the southern and western Pampas ecoregion [60]. Furthermore, as temperatures rise, certain crops may become more favorable than others. For instance, soybean crops might be more favored than corn and wheat crops [60]. The possible rodent community adaptations and dynamics that might arise from these changes should be studied as they may provide further insights into the potential distribution of *C. musculinus* and AHF.

While interpreting the inferred range shifts in rodent distribution from our models, we should consider some limitations. First, ecological systems and interactions between species and the environment are complex; some species might exhibit adaptive responses that models may not capture or might not be able to reach the predicted suitable areas due to natural or manmade barriers. For example, the fragmentation of ecosystems can create barriers for dispersal and migration, hindering the ability of species to move to suitable habitats in response to climate change [61, 62].

Secondly, due to limited availability, the data used to train the models had a temporal mismatch. Updated species occurrence data that closely aligns with the temporal resolution of bioclimatic and landscape variables could strengthen the reliability of the findings. Another limitation of this study is the lack of additional environmental factors. Incorporating other variables (which are currently unavailable for years under study) that may influence the distribution of the species could offer more precise predictions. Pardiñas et al. [63] suggested that rivers and their surrounding mesic conditions could have functioned as corridors for *C. musculinus* after the establishment of agricultural settlements in Patagonian River valleys, allowing the species to increase in abundance. Hence, future studies could incorporate distance to water bodies to account for this phenomenon. Additionally, including types of cultivated cropland could be beneficial. Previous studies have shown that corn crops and their stubbles are positive predictors of *C. musculinus* abundance [19]. Therefore, incorporating crop type data could help enhance model predictions. Our study relied on open-source data availability. Collaborations with governmental or private entities that control the data sources listed above will benefit the predictions generated by our models. Finally, considering species interactions through conducting multispecies modeling approaches could improve the predictions, as the presence of other species can influence the population dynamics and distribution of *C. musculinus* [64, 58].

Addressing the challenges of *C. musculinus* shifts requires the consideration of possible interventions and their effectiveness. Implementing public education campaigns can enhance awareness about JUNV transmission and promote preventative behaviors. Moreover, training health professionals on AHF diagnosis, treatment and prevention and organizing immunization campaigns with the Candid#1 vaccine are crucial. The assessment of pre- and post-campaign surveys and analysis of infection rates could be a tool to ensure these interventions effectively reduce the risk of transmission.

## Conclusions

Our study provides a valuable tool for understanding general trends in the distribution of *C. musculinus* response to climate change. Knowing the potential areas of habitat suitability for this rodent and the factors that could be shaping these shifts, provides information for public health interventions targeting AHF prevention. Preventive measures such as rodent ecological studies and epidemiological surveillance could be directed to regions with potential expansion of *C. musculinus*. Moreover, human vaccination efforts, which have been incorporated into the National Immunization Calendar in the endemic area of the disease since 2007 [65], could also be enhanced to address the risk of shifts in the disease dynamics, particularly in the identified hotspots for potential disease transmission. Our findings underscore the role of modeling predictive tools in guiding public health strategies, emphasizing the crucial need for continuous innovation and collaboration to effectively address evolving disease dynamics amidst the challenges posed by climate change.

## Acknowledgments

We thank Janet Foley and Isabel Fletcher for providing useful comments and feedback on the manuscript.

## Supporting information

**S1 Table. Bioclimatic and landscape variables relevant to *C. musculinus* distribution.**

**S1 Fig. Current registered distribution of *C. musculinus* in Argentina’s ecoregions**. Black dots represent occurrences of *C. musculinus* from 1990 to 2021 across ecoregions in Argentina, based on distribution data retrieved from the Global Biodiversity Information Facility (GBIF).

**S2 Fig. Boxplot of bioclimatic and landscape variable importance across machine learning classifiers.** Each boxplot summarizes the distribution of the 1000 simulation variable importance results for machine learning classifiers: Random Forest (RF), Extra Trees (ET), Extreme Gradient Boosting (XGB), and Light Gradient-Boosting (LGBM).

**S3 Fig. Correlation heatmap between bioclimatic and landscape variables**. The Pearson correlation coefficient was used to measure the correlation between bioclimatic and landscape variables. Only one variable from each pair of highly correlated variables (≥ 0.7) was included in the species distribution models.

**S4 Fig. Predicted distribution of *C. musculinus* under climate change scenarios**. The potential distribution of the rodent under two climate change scenarios is shown in Argentina’s ecoregions. The maps show distribution probability under a) RCP4.5 scenario for year 2050, b) RCP4.5 scenario for year 2070, c) RCP8.5 scenario for year 2050, and d) RCP8.5 scenario for year 2070. Note: Higher values and darker blue colors indicate a greater likelihood of *C. musculinus* habitat suitability.

**S5 Fig. Partial dependence of *C. musculinus* distribution on bioclimatic and landscape variables**. a) Secondary land (percentages of secondary coverage), b) Mean temperature of driest quarter (°C), c) Isothermality (percentage of temperature variability), d) Primary land (percentages of primary coverage), e) Elevation (meters), f) Cropland (percentages of cropland coverage). Note: The color of the lines indicates the machine learning classifier used: Random Forest (RF) in blue, Extra Trees (ET) in orange, Extreme Gradient Boosting (XGB) in green and Light Gradient-Boosting (LGBM) in red.

